# Identification of a female determinant gene for the sexual determination of a hemipteran insect, the brown planthopper

**DOI:** 10.1101/775577

**Authors:** Ji-Chong Zhuo, Hou-Hong Zhang, Yu-Cheng Xie, Han-Jing Li, Qing-Ling Hu, Chuan-Xi Zhang

## Abstract

The sex determination mechanism for hemipteran species remains poorly understood. During the sex determination of the brown planthopper (BPH), *Nilaparvata lugens*, one species of Hemiptera, the functions of *doublesex* (*Nldsx*) and *NlTra-2* (*NlTra-2*) genes were identified in our previous studies. Here, we identify an upstream gene for *Nldsx* in the sex determination cascade, *NlFmd*, which acts as female determinant gene for *N. lugens*. The sex-specific transcript of *NlFmd* (*NlFmd-F*) encodes an arginine/serine-, and proline-rich protein that is essential for female development. The knockdown of *NlFmd* resulted in the development of pseudomales, with sex-specific alternative *Nldsx* processing, and maternal RNA interference (RNAi) against *NlFmd* generates male-only progeny. Moreover, homologous genes for *NlFmd* have also been identified in two rice planthopper species, the white-backed planthopper (WBPH, *Sogotalla furcifera*) and the small brown planthopper (SBPH, *Laodelphax striatellus*), and these genes appear to be involved in the sex determination cascades for these species. Our data suggest that the sex determination cascade in Delphacidae is conserved.

## Introduction

Insects have evolved an astonishing variety of molecular mechanisms to achieve sexual differentiation, which is a fundamental characteristic of life. Although *doublesex* (*dsx*) has been identified as the most well-conserved gene at bottom of the sex determination cascade, the identified upstream regulators have demonstrated the broad diversity of regulators among different insect species (Shukla and Nagaraju, 2010). For example, the primary signals for sex etermination in *Drosophila melanogaster*, the honeybee, the silkworm, and mosquitoes appear to be quite different and extremely variable. In *D. melanogaster*, the direct actions of several X-encoded signal element (XSE) proteins determine the female or male gender (Cmkovich et al., 2007). In the mosquito, *Aedes aegypti*, a transformer 2-like gene, *Nix*, serves as the primary gender determining signal (Hall et al., 2015). In *Apis mellifera*, a hymenopteran species, the gene *csd* acts as the primary signal according to its allelic composition (Beye et al., 2003). In *Bombyx mori*, a single female-specific piwi-interacting RNA (piRNA) is the primary sex determinant (Xu et al., 2017).

In the sex determination cascade, proteins that directly regulate the alternative RNA splicing of *dsx* have only been identified in a few insect species. In female *D. melanogaster*, the female-specific genes *transformer* (*DmTra*) and *transformer-2* (*DmTra-2*) regulate the alternative RNA splicing of *dsx* pre-mRNA (Belote and Baker, 1982; Hoshijima et al., 1991; McKeown et al., 1987). The DmTra protein belongs to serine-arginine (SR) protein families but lacks an RNA-recognition motif (RRM), which is essential for the binding of pre-mRNA (Hedley and Maniatis, 1991). DmTra requires another protein, Transformer-2 (*DmTra-2*), which provides the RRM domain that binds the purine-rich enhancer elements in *dsx* pre-mRNA, resulting in the female-specific splicing of *Dmdsx*, in which exon 4 is retained (Hedley and Maniatis, 1991; Hoshijima et al., 1991). In male *Drosophila*, male-type *Dmdsx* is produced by default splicing. In *Apis mellifera*, the gene *fem*, which encodes an SR-type protein, functions similarly to *DmTra* during sex determination (Hasselmann et al., 2008). In *B. mori*, the female splicing of *Bmdsx* pre-mRNA represents the default mode, and the female exon lacks putative binding sites for Tra and Tra-2. Moreover, no *tra* homolog has been identified in *B. mori*, and the gene *Bmtra-2* is not involved in the regulation of sex-specific *Bmdsx* pre-mRNA splicing (Niu et al., 2005; Suzuki et al., 2001; Suzuki et al., 2012). Therefore, the molecular mechanism through which the sex-specific splicing of *Bmdsx* pre-mRNA occurs in *B. mori* appears to be very different from the mechanism that regulates Diptera *dsx*.

In our previous studies, both *Nldsx* and *Nltra2* were found to play important roles during the sex determination in the brown planthopper (BPH, *Nilaparvata lugens);* moreover, the sex-specific splicing of *Nldsx* is common, with *Dmdsx* exhibiting different repeat nucleotide sequences on the female-specific exon (Zhuo et al., 2018; Zhuo et al., 2017). Although no homologous *tra* gene has been reported in BPH, we speculate that such a gene exists in BPH that plays a function similar to that of *tra*. In this study, we sequenced the early embryonic transcriptome and chose 350 genes for a large-scale functional screen using RNA inference (RNAi). We identified a new gene involved in the sex determination cascade of BPH, named *female determinant factor* (*NlFmd*). Moreover, we also identified *NlFmd* homologs that play roles during the sex determination of the white-backed planthopper (WBPH, *Sogotalla furcifera*), and the small brown planthopper (SBPH, *Laodelphax striatellus*), two other planthoppers in the Delphacidae family, suggesting that the direct upstream genes of the sex determination cascade might be conserved in Delphacidae.

## Materials and Methods

### Insect rearing

The BPH populations used in this study were collected in Hangzhou (30°16’9N, 12°11’E), China, in 2008. BPHs were reared at 26 ± 0.5°C, on rice seedlings (Xiushui 128), under a 16 hr light:8 hr dark photoperiod.

### Sequence analysis and alignments

The Splign tool from NCBI (www.ncbi.nlm.nih.gov/sutils/splign/splign.cgi) was used to predict *NlFmd* exons and introns by aligning the sequences of the reverse transcriptase polymerase chain reaction (RT-PCR) products with the genomic sequence of BPH (PRJNA177647). The alignment of homologous *Fmd* gene sequences was generated using Clustal X (JEANMOUGIN et al.1998) and GENEDOC (Nicholas and Nicholas, 1997, http://www.softpedia.com/get/Science-CAD/GeneDoc.shtml).

### RNA interference (RNAi)

Two regions that were common among the *NlFmd* isoforms were used as DNA templates for double-stranded RNA (dsRNA) synthesis. The MEGAscript T7 High Yield Transcription Kit (catalog no.AM1334; Ambion) was used to synthesize the dsRNAs, according to the manufacturer’s instructions. The concentration of the product was quantified using a NanoDrop 2000 (Thermo Fisher Scientific, Waltham, MA). Afterward, we followed the BPH dsRNA treatment method, as described by Xue et al. (Xue et al. 2015, http://dx.doi.org/10.1038/protex.2015.005).

### PCR of the male-specific genomic DNA fragment

Genomic DNA for genotyping PCR was purified from each treated planthopper using the Wizard GenomicDNA Purification Kit, according to the manufacturer’s instructions (Promega, Madison, WI), starting with 600 ml Nuclei Lysis Solution. The genomic DNA was concentrated and desalted by isopropanol precipitation. The DNA concentrations were measured and diluted to 50 ng/ml. Primers for a male-specific sequence (PM3nF: 59-GAGCTGGAGTGTGTT GATGT-39 and PM3nR: 59-CAAGTTGATTGAAACAGACT-39) and primers for a sequence common to both sexes (PM3femaleF: 59-ATCATACCCGCATGTTGGAC-39 and 59-GAATCTGAAGGGAGCTGTGC-39) were used, as previously described (Kobayashi and Noda 2007). PCR was performed on 1 ml genomic DNA in a 25-ml reaction mixture (50 units/ml Taq DNA polymerase, W/Mg^2+^ buffer for DNA polymerase [Biocolors, Shanghai, China]; 2 mM dNTPs, [TaKaRa]). Thirty cycles of amplification were performed, each of which consisted of denaturation for 30 sec at 94°C, annealing for 30 sec at 56°C, and extension for 30 sec at 70°C. Samples of the PCR reactions were analyzed on 2% agarose gels.

### Quantitative real-time PCR (qRT-PCR) analysis

Total RNA was first isolated from BPHs and tissues using RNAiso Plus (TaKaRa), and then each RNA sample (1 μg) was reverse transcribed using the Prime Script first-strand cDNA synthesis kit (catalog no. 6110A; TaKaRa). The internal control for qRT-PCR experiments was the 18S rRNA gene of BPH. The SYBR Premix ExTaq Kit (TaKaRa) was used for qRT-PCR. The relative quantitative method was used to evaluate quantitative variations.

### Semiquantitative RT- PCR

Total RNA was extracted from embryos, first-, second-, third-, fourth-, fifth-instar nymphs, and female and male adults. A total of 1 mg RNA was used to perform reverse transcription in 20-ml reactions, using Quant reverse transcriptase (TIANGEN, Beijing, China) according to the manufacturer’s instructions. Reverse transcription reactions were then diluted 10 times, and 1 ml was used in the following PCR reaction.

### Dissection and fertility analysis

Third-instar nymphs were injected with ds*GFP* or ds*NlFmd*, and then either dissected or used for fertility analysis 3 days after emergence. The treated BPH pairs were reared on rice seedlings, and the numbers of offspring and eggs were examined 10 days after deposition. Each pair of adults were reared on the same type of rice seedling. At least 10 pairs of adults per treatment were tested.

### Image processing

dsRNAs targeting *GFP* and the region of *Fmd* were injected into third-instar nymphs, and phenotypes were determined after emergence, using a DFC320 digital camera attached to a LEICA S8APO stereomicroscope.

### Maternal RNAi and offspring counting

Virgin female adults were collected 12 hr after emergence, injected with ds*GFP* and ds*NlFmd*, reared for 3 days, and then mated with wild-type males. Each pair was reared separately. The offspring were reared, and the numbers of each sex were counted. Dead eggs were dissected and counted from the seedlings after the larva failed to hatch for three consecutive days.

## Results

### The sequences and expression of *NlFmd* in BPH

*NlFmd* was identified by using large-scale RNAi screens of *N. lugens* embryonic transcriptomes. Two *NlFmd* splicing transcripts were characterized by comparison with BPH genome sequences (Bioproject PRJNA177647). We found that both the exon skipping and 5’ splice sites resulted in the two primary splicing isoforms of *NlFmd* sharing the same 5’ untranslated region (UTR); however, these isoforms differed in their downstream exon compositions (Fig. 1A). In this study, a pair of primers were designed to test the expression levels of the two primary splicing transcripts, and the semi-quantitative PCR results showed that the transcript without exon 6 (150 bp) was female-specific (*NlFmd-F*), whereas the transcript containing exon 6 was expressed in both females and males (*NlFmd-C*) (Fig. 1B). Moreover, the knock-down of exon 14 in *NlFmd* transcripts was able to silence female-specific *NlFmd*, indicating that exon 14, which contains a stop codon, is present in the female-specific transcript.

**Fig. 1.**
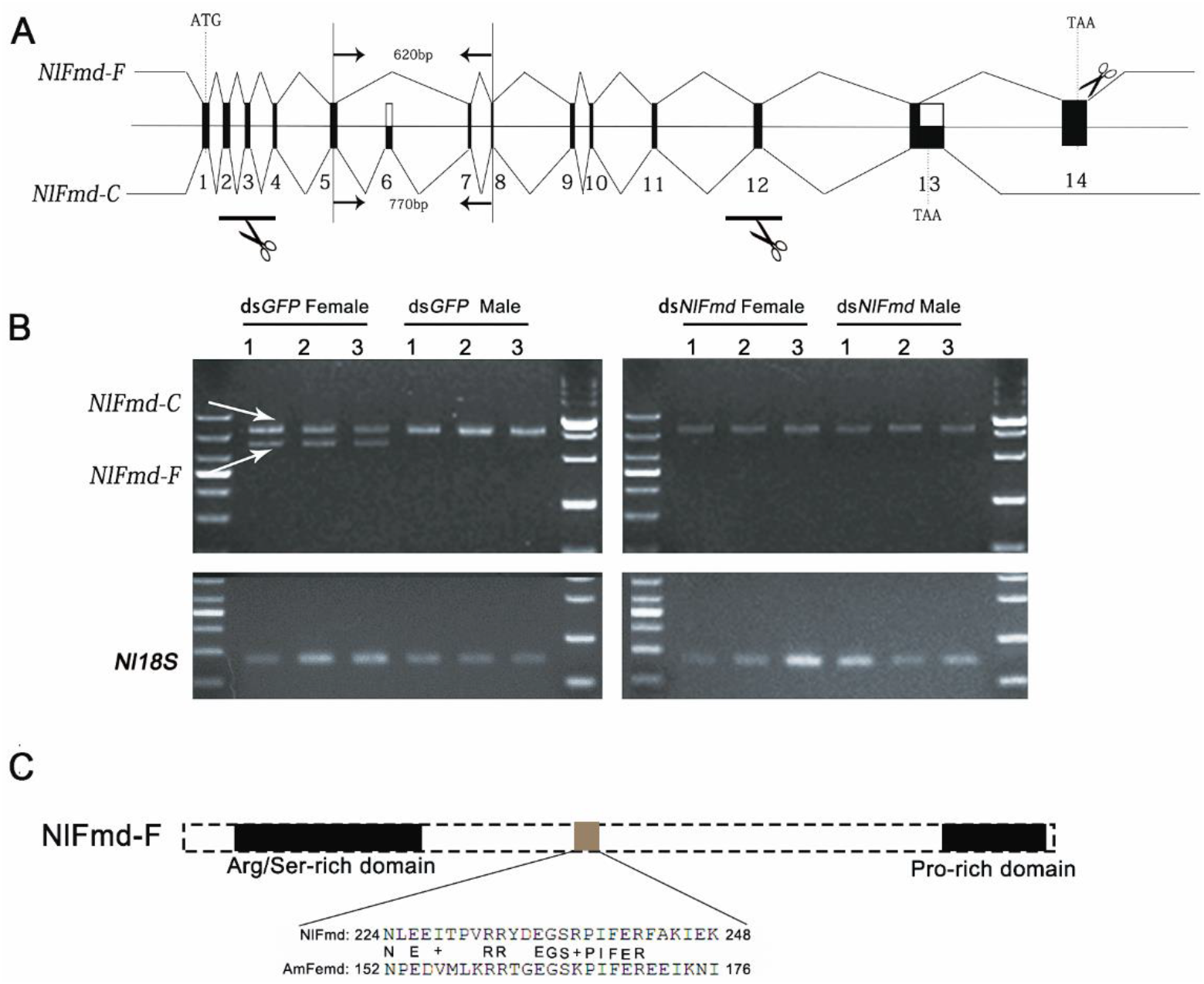
Alternative splicing and sequence features of *NlFmd*. A) The two primary alternative splicing isoforms of *NlFmd*. B) The differential expressions of *NlFmd-F* and *NlFmd-C* in males and females. C) Motifs and structures of *NlFmd-F*. Scissors in A) represent the RNAi target sequences used in this study.

The female-specific *NlFmd* encodes a 613-amino-acid protein, which contains Arg/Ser- and Pro-rich domains and is a member of the SR-type proteins, with no similarity to other proteins and no obvious motifs identified in other proteins (Fig. 1C). The protein translated by the non-sex-specific *NlFmd-C* also belongs to the SR-type family of proteins. When the NlFmd protein sequence was compared with that for Tra from *D. melanogaster* and with that for Fem from *A. mellifera*, only a 27-amino-acid region with low similarity to AmFem was identified, and no similarity with *D. melanogaster* Tra was found. Thus, we name the gene *Fmd* in planthoppers, instead of *Tra* (gray box in Fig. 1C).

### *NlFmd* influences the soma development and fertility of female BPHs

To study the functions of *NlFmd*, RNAi-mediated gene silencing was performed, using dsRNAs designed to target three different regions of *NlFmd* transcripts (Fig. 1A). Specifically, RNAi knockdown of *NlFmd* in third-instar BPH nymphs resulted in females that developed into pseudomales, with a series of masculine phenotypes, such as shorter ovipositors, male-specific claspers, and intromittent organs, and the females lost their fertility, with undeveloped ovaries (Figs. 2 and 3; Table 1). However, when the non-sex-specific *NlFmd-C* was targeted with RNAi in male BPHs, no difference was found between ds*NlFmd-* treated males and control males, from somatic development to fertility, and the knockdown of exon 14 (*NlFmd–F*) also showed no influence in males (Figs. 2B and 3C; Table 1).

**Fig. 2.**
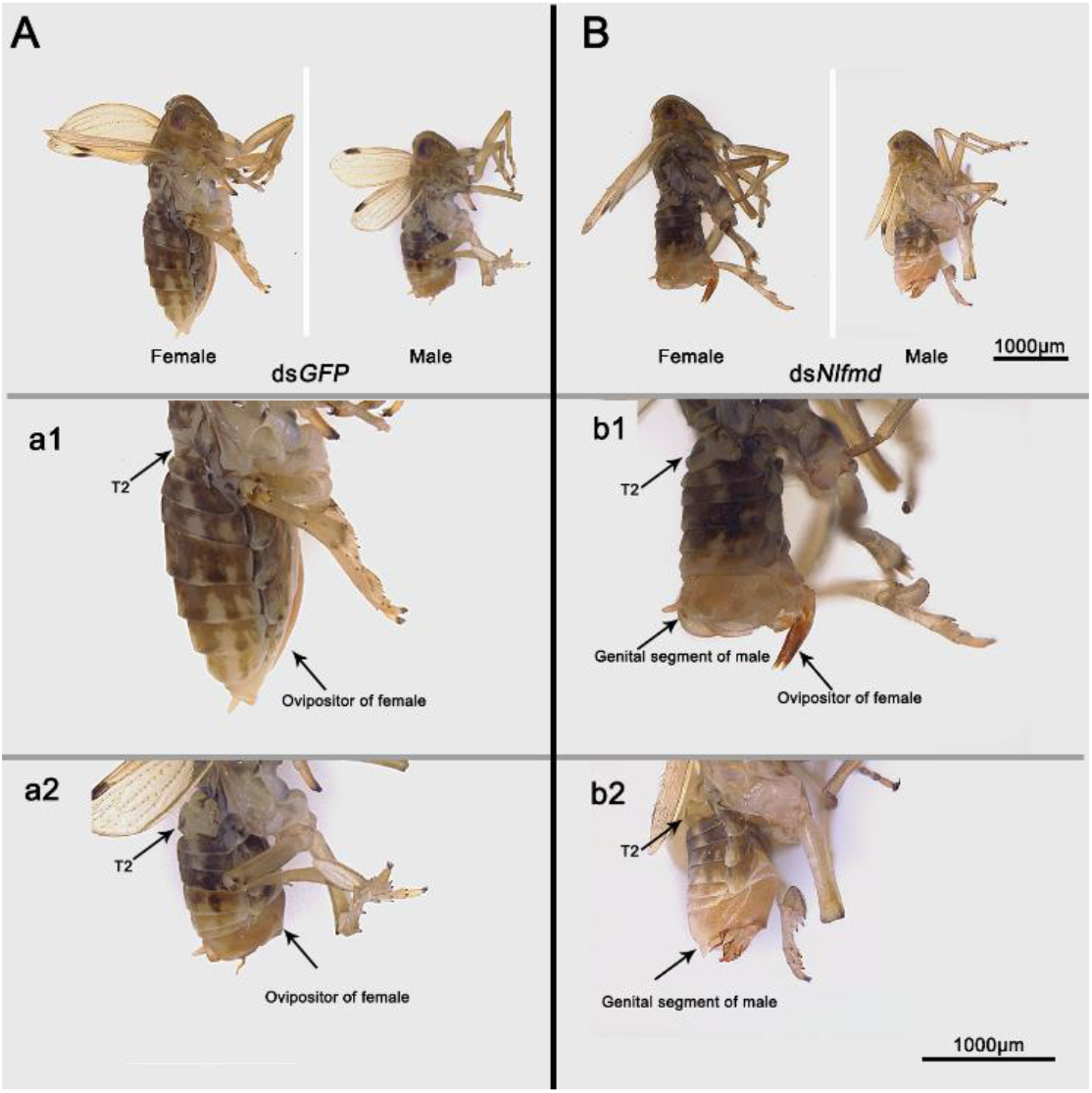
The soma development of ds*GFP*- and ds*NlFmd*-treated BPHs. A) The soma development of ds*GFP*-treated BPH females and males. B) The soma development of ds*NlFmd*-treated BPH females and males. a1, a2, b1, and b2 show the abdomen, and T2 indicates the 2^nd^ tergite in the abdomen.

**Fig. 3.**
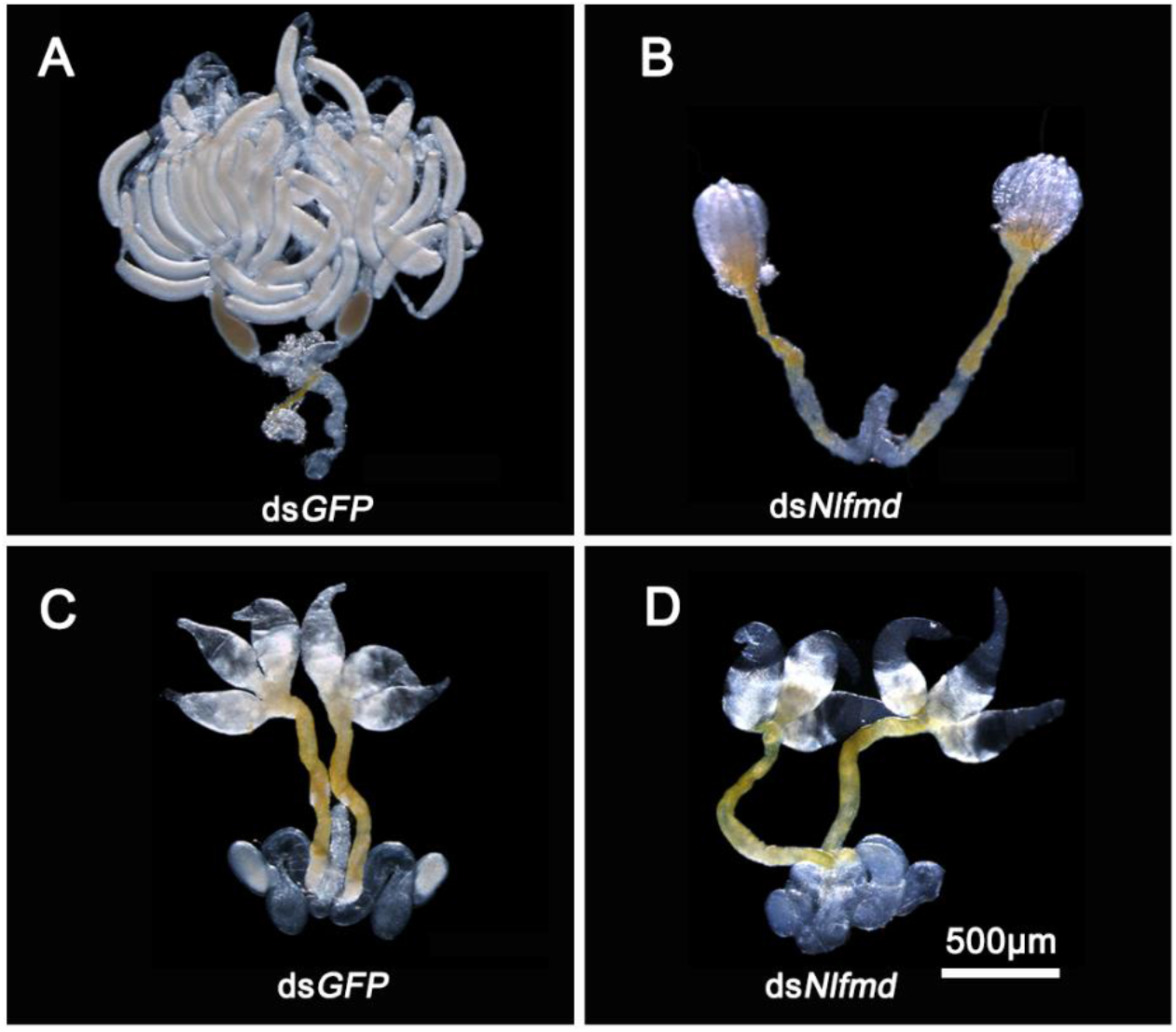
The development of ovaries and testes. A and C) The ovaries and testes of ds*GFP*-treated BPHs. B and D) the ovaries and testes of ds*NlFmd*-treated BPHs.

**Table1.** Influence of *NlFmd* on female and male fertility (offspring number)

| Treatment | #1 | #2 | #3 | Average |
| --- | --- | --- | --- | --- |
| **dsGFP♀ X dsGFP ♂** | 238 | 134 | 208 | 193±38 |
| **dsNIFmd♀ XdsGFP ♂** | 0 | 0 | 0 | 0 |
| **dsGFP♀ X dsNIFmd ♂** | 155 | 206 | 200 | 187±20 |

### *NlFmd-F* regulates *Nldsx* alternative splicing

The *dsx* gene functions as a stable-base molecular switch, and the differential splicing of this gene results in the development of sexually dimorphic traits. In our previous study, *dsx* in *N. lugens* (*Nldsx*) was found to have two sex-specific isoforms: female-specific *Nldsx^F^* and male-specific *Nldsx^M^. Nldsx* controls sexual dimorphism based on male-specific expression, whereas female sexual development appears to occur by default. In this study, we found that ds*NlFmd*-treated females developed into pseudomales. Based on the results of qRT-PCR, we confirmed that ds*NlFmd*-treated females produced male-specific *Nldsx^M^* at a similar level to that produced by ds*GFP*-treated males, and the amount of *Nldsx^F^* decreased by more than 80 percent compared with that in ds*GFP*-treated females, indicating that NlFmd is upstream of *Nldsx* (Fig. 4).

**Fig. 4.**
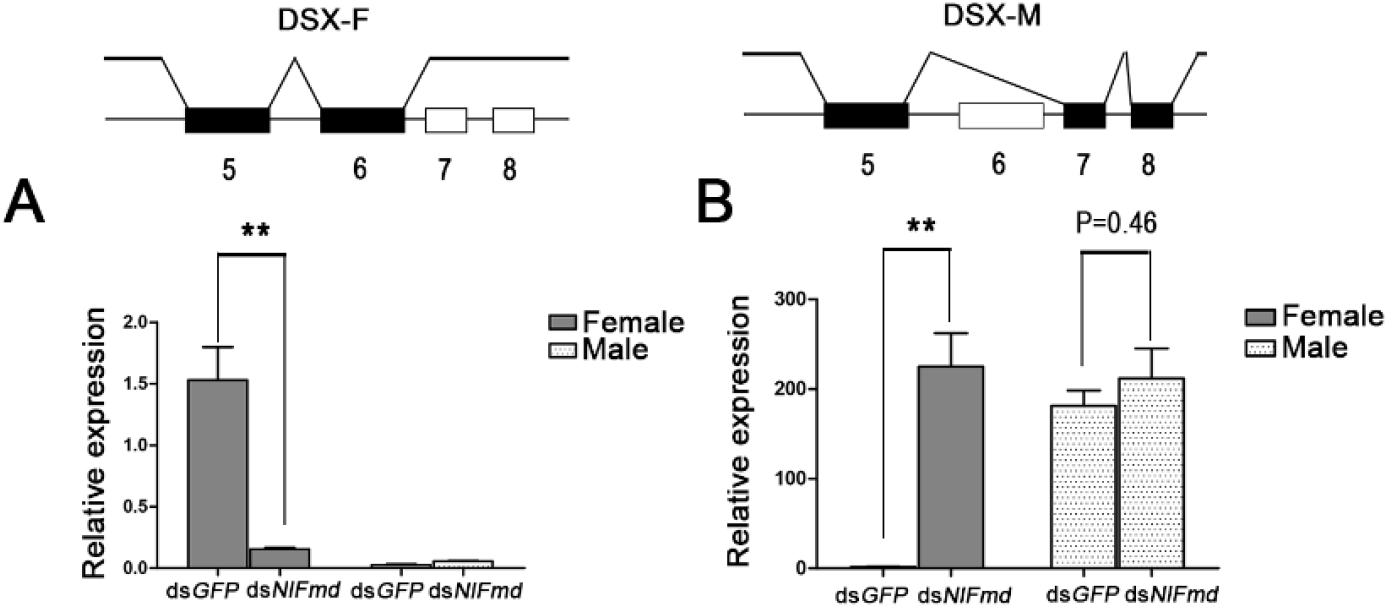
The *NlFmd*-mediated regulation of sex-specific *Nldsx* expression in females. A) The knockdown of *NlFmd* decreased the female-specific expression of *Nldsx*. B) The knockdown of *NlFmd* increased the expression of the male-specific *Nldsx* isoform in females.

### *NlFmd* is required for female embryo development

During embryo development, sex is established during the early stages, and in BPH, female-specific *Nldsx* transcripts could be detected starting at 60 hr, which indicates that sex was established prior to this time point. *NlFmd-C* and *NlFmd-F* transcripts could be detected by RT-PCR in the newly laid eggs and during all stages, suggesting that both genes were maternally inherited (Fig. 5A). However, the expression of *NlFmd-F* decreased sharply during the early period of the embryo stage, from 0-12 hr, and then increased starting at 24 hr before finally reaching a normal and stable expression level starting at 48 hr, which corresponded to the transcription of *Nldsx* (Fig. 5A and B). When we used ds*NlFmd* to knock down the expression of *NlFmd* in newly emerged females, the number of offspring produced by ds*NlFmd*-treated females was only 10 percent that produced by ds*GFP*-treated females; moreover, the offspring of ds*NlFmd*-treated females were all males, with a hatching rate of approximately 50 percent (Fig. 5C and D). To further confirm the sex of these insects, genomic DNA was isolated for PCR verification, using the male-specific primers PM3n (Kobayashi and Noda, 2007). Because *NlFmd* is essential for female development, the female embryos likely died without functional *NlFmd*, resulting in the male-only offspring. We further tested the fertility of these male offspring and found no differences compared with control males (data not shown).

**Fig. 5.**
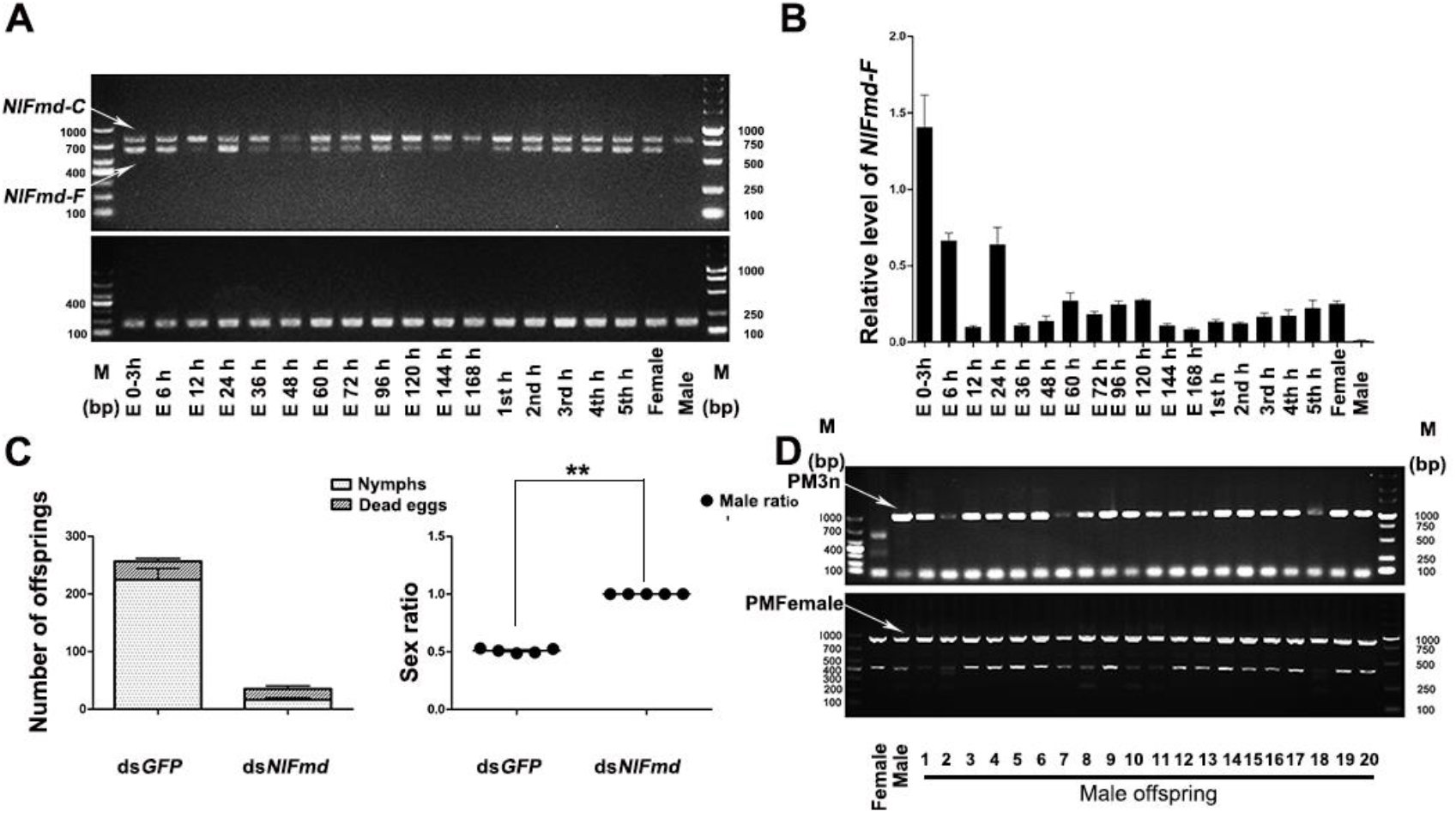
The influence of maternal RNAi *NlFmd* knockdown. A) The expression of *NlFmd* during different development stages in both sexes. B) The expression of *NlFmd-F* during different development stages in both sexes, assessed by qRT-PCR. C) The effects of maternal ds*NlFmd* treatment on the embryonic development of offspring. D) The genotypes of offspring from *dsNlFmd*-treated females. The male-specific primers PM3n were used to identify male genotypes, whereas the non-sex-specific primers PMFemale were used as positive controls.

### *NlFmd* homologous genes in two other rice planthoppers

Similar to the BPH, both *S. furcifera* and *L. striatellus* are also destructive hemipteran pests found in rice ecosystems in Asia. We wondered whether the *NlFmd* homologs in these two delphacids had the same functions during sex determination.

We used the protein sequence for NlFmd to BLAST the transcriptomes of *S. furcifera* and *L. striatellus* and identified two *NlFmd* homologous genes, *SfFmd* and *LsFmd*, respectively. Based on multiple alignments of NlFmd, SfFmd, and LsFmd-F, we found that they shared a high degree of sequence similarity (Fig. 6), including the Pro/Phe-rich C-terminal domain, which is only in NlFmd-F, but not in NlFmd-C, which we believe to be essential for the function of *NlFmd* during sex determination.

**Fig. 6.**
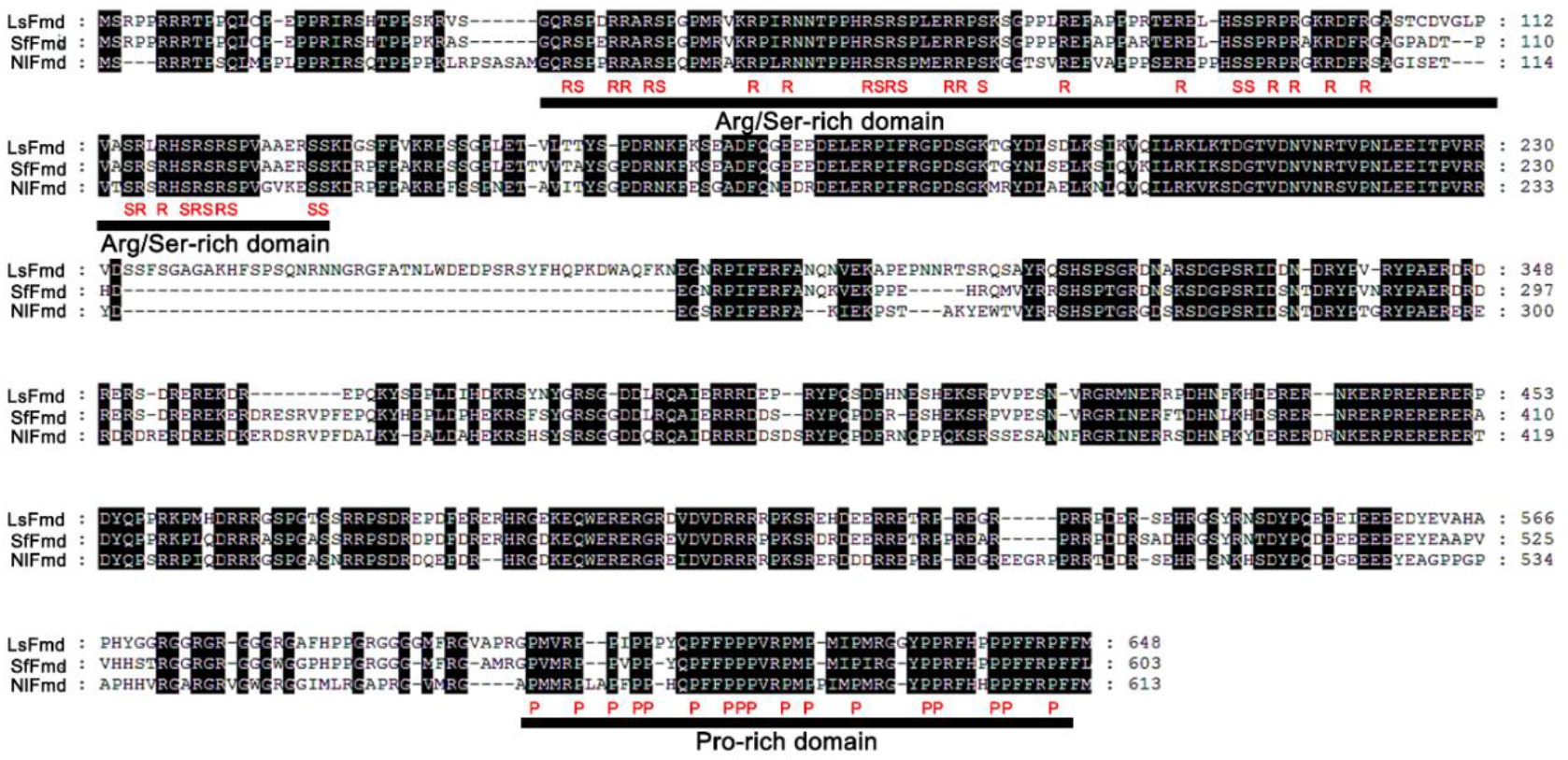
Alignments among the *Fmd* homologous sequences from three planthoppers. The Arg/Ser-rich domain and Pro-rich domain were indicated by a black lines.

We also knocked down the expression of *SfFmd* and *LsFmd* with RNAi in third-instar nymphs. After adult eclosion, we found that *dsSfFmd-* and ds*LsFmd*-treated females developed into pseudomales containing male external genitalia, similar to the phenotype observed for ds*NlFmd*-treated BPH females, whereas treated males showed normal soma development (Fig. 7). Together, these results suggested that *Fmd* might play a conserved function in these three delphacid species.

**Fig. 7.**
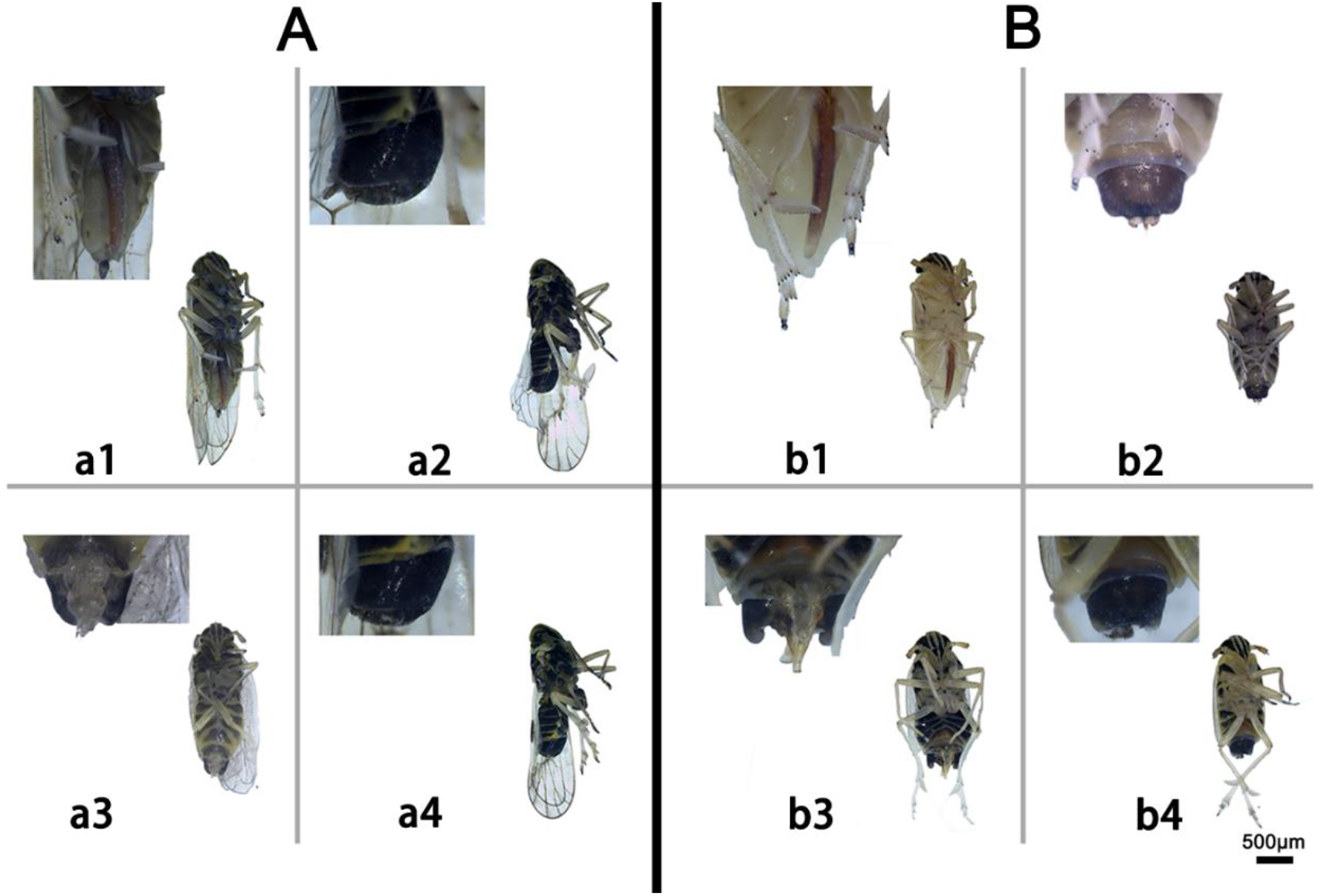
The influence of *Fmd* homologous genes on soma development in *S. furcifera* and *L. striatellus*. A) The phenotypes of *S. furcifera* adults after injection with ds*GFP* and *dsSfFmd*. a1 and a2, ds*GFP*-treated females and males, respectively; a3 and a4, dsSfFmd-treated females and males, respectively. B) The phenotype of *L. striatellus* adults after injection with ds*GFP* and ds*LsFmd*. a1 and a2, ds*GFP*-treated females and males, respectively; a3 and a4, ds*LsFmd*-treated females and males, respectively.

## Discussion

In this study, we characterized the gene *NlFmd* as an upstream regulator of *Nldsx* in the BPH sex-determination pathway. *NlFmd* has equivalent functions as *AmFem* and *DmTra*, which regulate *Nldsx* pre-mRNA through female-specific splicing isoforms. The knockdown of *NlFmd* in females reverses the sex-specific expression of *Nldsx*, with the female-specific expression of *Nldsx^F^* reducing greatly and the male-specific *Nldsx^M^* being expressed at similar levels as in control males, resulting in the development of pseudomales. Moreover, the knockdown of *NlFmd* homologous genes in two other delphacid species also resulted in the development of females into pseudomales. Therefore, *Fmd* might have conserved functions during sex determination in Delphacidae.

The maternal input of *tra* mRNA or protein into eggs has been demonstrated for many insect species(Verhulst et al., 2010), and a similar phenomenon has been described for *NlFmd* in BPH. During the 192-hr long BPH embryonic stage, the mRNA expression of functional, female-specific *NlFmd-F* sharply decreased from a very high level to a low level during the first 12 hr, increased again at 24 hr, and then maintained a relatively lower expression level starting at 48 hr until the adult stage. Among the two primary alternative splice forms of *NlFmd*, exon 6 is non-sex-specific, and alternative 5’ splicing in exon 13 results in different C’-protein sequences between NlFmd-F and NlFmd-C, which is believed to be the most important domain for its functions. Because no male-specific isoform has been identified in males, the non-sex-specific alternative isoform *NlFmd-C* may represent the default mode, and we believe that a female-specific factor may be involved in the production of the female-specific alternative transcript *NlFmd-F*.

The alternative splicing cascade represents the basic strategy for insect sex determination. In *Drosophila, DmTra-2* is essential for the alternative regulation of *Dmdsx* because it provides the RRM domain that is necessary for the recognition of the ESE motif in *Dmdsx* exon 4 that promotes the female-specific splicing of *Dmdsx*. In our previous study, a *tra-2* homolog in BPH (*NlTra-2*) was found to be required for the sex determination of BPHs. Whether and how *NlTra-2* cooperates with *NlFmd* to regulate the alternative splicing of *dsx* pre-mRNA requires future study.

In rice ecosystems in Asia, rice planthoppers, including BPH (*N. lugens), S. furcifera*, and *L. striatellus*, have become destructive insect pests, the effective and biologically-based sterile insect techniques (SITs) represent promising control methods. Because *Fmd* is a key sex determination gene in these three rice planthoppers, we hope these findings form the basis for a new method for controlling these agriculture pests.

## Acknowledgements

This work was supported by the National Natural Science Foundation of China (grant no. 31871954 and 31630057).

